# Comment on “Connexins evolved after early chordates lost innexin diversity”

**DOI:** 10.1101/2023.01.05.522624

**Authors:** Ilya V. Kelmanson, Evgeny A. Bakin, Izhar Karbat, Georgy A. Slivko-Koltchik, Eitan Reuveny, Yuri V. Panchin

**Affiliations:** Department of Biomolecular Sciences, Weizmann Institute of Science, Rehovot, Israel; Bioinformatics Institute, Saint-Petersburg, Russia; Kharkevich Institute for Information Transmission Problems, Russian Academy of Sciences, Moscow, Russian Federation; A.N. Belozersky Institute of Physico-Chemical Biology, Moscow State University, Moscow, Russian Federation

## Abstract

Gap junctional (GJ) intercellular channels are an important way of intercellular communication. Currently, two unrelated families of GJ proteins, innexins/pannexins, and connexins, are known, either or both of which are present in most multicellular animals. A striking exception is the echinoderms which have functional GJs but until recently were believed to be lacking both innexins and connexins, which suggests the presence of the third, yet the unknown family of GJ proteins. In the recent work Welzel and Schuster (Welzel and Schuster, 2022) have reported several putative innexins and one connexin from echinoderms, therefore undermining such a hypothesis. Here we provide evidence showing that all reported connexin and innexin sequences from echinoderms are cross-species contaminations, indicating that a search for a third GJ protein family is still a subject of immediate scientific research interest.

## Introduction

Gap junctional intercellular channels directly connect the cytoplasm of adjacent cells in multicellular animals, allowing the passage of ions, small molecules, and membrane potential changes. GJs participate in several functions and are found in virtually all solid animal tissues (Goodenough and Paul, 2009). So far there are no taxonomic groups among Metazoa for which the absence of functional intercellular channels is decisively shown. GJ channels are formed by proteins belonging to two unrelated protein families, connexins and innexins. Despite the lack of sequence homology between them, the topology and functions of innexin and connexin intercellular channels are remarkably similar (Skerrett and Williams, 2017). Connexins are found only in vertebrates and their close relatives, tunicates. Innexins are present in most taxons of invertebrate animals. Vertebrates and tunicates also have proteins homologous to innexins called pannexins (Panchin et al., 2000; Baranova et al., 2004), however, either most or even all of them have lost the ability to form intercellular channels.

With the growing access to high-throughput sequencing, the vast amount of genomic information for all kinds of living matter has become available for analysis, allowing not only to confirm the presence of a particular gene family in a certain organism or a taxon but also to prove the absence of it. The earlier analysis of the innexins/pannexins and connexins distribution (Slivko-Koltchik et al., 2019) has revealed several clades of multicellular animals lacking both gene families. One of those clades is Echinodermata – a large phylum that includes well-studied animals such as sea urchins and sea stars. Two possible conclusions can be drawn from this result: either some animals do not use intercellular channels or there are intercellular channels built of some other proteins, neither innexins nor connexins. Several independent studies showing the GJ-like structures and electrical coupling between the blastomeres of echinoderm embryos (Tupper et al., 1970; Yazaki et al., 1999; Slivko-Koltchik et al., 2019) support the latter assumption.

If found, the new GJ proteins would shed light on many basic aspects of intercellular channel biology. However, there is an additional intriguing possibility. This family might appear to be unique for echinoderms like connexins are unique for chordates, but it also might be more universal. The members of this family in other phyla (including the chordates, who are the closest relatives of echinoderms) might also form intercellular channels or they may have other important functions, similar to vertebrate pannexins that form large pore ion channels permeable for ATP. This makes searches for putative echinoderm GJ proteins a highly relevant scientific task.

The recent publication of Welzel and Schuster (Welzel and Schuster, 2022) has challenged this view. In the course of screening for glycosylation sites in GJ proteins in the public genome, transcriptome, and protein databases, they have identified several innexins and one connexin sequences in echinoderms. If true, such findings perfectly explain the electrical connections in the echinoderm embryos and make the speculations regarding the third GJ protein family unnecessary and redundant. Since this question has important biological implications, we have performed an additional analysis of the sequences found in the discussed work. Our results indicate that all the GJ genes attributed to echinoderms are cross-species contaminations from other animal phyla.

## Methods

BLASTN and BLASTX searches were performed against NCBI sequence databases. To compare the presence of putative innexin\pannexin-like sequences in echinoderm TSAs to TSA projects from other taxonomic groups, we have selected 5 protein sequences of innexins and pannexins from different taxonomic groups (NP_001274699.1 innexin inx1-like [Hydra vulgaris], JAP56454.1 hypothetical protein TR161073 [Schistocephalus solidus], TRY68384.1 hypothetical protein TCAL_01429, partial [Tigriopus californicus], NP_001309635.1 Innexin [Caenorhabditis elegans], NP_001002005.2 pannexin-2 [Mus musculus]) and concatenated them into one >inn sequence (see Supplementary materials). This sequence was run with TBLASTN with a threshold of E-value <1e-10 against TSA databases of Echinodermata, Porifera, Fungi, Chlorophyta, Rhodophyta, Phaeophyceae, Platyhelminthes, Nematoda, Annelida, Copepoda, Gastropoda, Bivalvia and Cephalopoda (only the TSAs containing more than 1000 sequences were taken into analysis). The taxons for comparison were selected so that the total number of available TSAs containing more than 1000 sequences fitted in the range from 20 to 200, which is comparable to that of echinoderms (66). The taxonomic ranks are different due to the unequal representation of different groups in the databases.

## Results and discussion

The huge speed of data accumulation in modern high-throughput sequencing projects makes the contamination of the databases with non-specific sequences inevitable. This especially concerns the sequences of samples collected from natural environments where different organisms can live in close relationship with each other and one bigger organism can be populated by thousands of smaller inhabitants. This factor must be taken into account when analyzing such data.

Welzel and Schuster reported 4 putative innexin\pannexin related sequences that were found in Transcriptome Shotgun Assembly (TSA) sequences databases of 4 echinoderm species (Ec_Aja_01 from Apostichopus japonicus; Ec_Apu_01 from Arbacia punctulata; Ec_Aru_01 from Asterias rubens; Ec_Pli_01 from Paracentrotus lividus) after the filtering procedure suggested in the paper. Those sequences were found in 6 out of 65 Echinodermata TSA assembles. At the same time, no innexin\pannexin related sequences were found in any of 35 assembled echinoderm genomes or 277160 NCBI annotated proteins or 3417970 nucleotide sequences.

In the case of Asterias rubens both innexin\pannexin related sequences GHKZ01096691.1 and GHKZ01082365.1 were found in one TSA (GHKZ01000000 Asterias rubens tube foot transcriptome; sequence BioProject: PRJNA268905, BioSample: SAMN03246764, Sequence Read Archive (SRA): SRR1685733). Thus the presence of innexins in Asterias rubens is only supported by sequences from a single SRA with 6 Gb reads out of 10 SRA experiments containing 362 Gb total. Accurately assembled Asterias rubens genome based on 15 related DNA SRA experiments or 24412 recorded proteins contain no reads related to putative innexin\pannexin sequences, including GHKZ01096691.1 and GHKZ01082365.1. Importantly, Asterias rubens putative innexin\pannexin related sequences are most similar to nematode proteins that are not expected to be related to echinoderms. Thus, it is most likely that a BioSample was contaminated with untargeted species, probably nematode. The origin of this sample - tube foot provides an easy way for contamination to occur.

The case of GHCH01013653.1 from Apostichopus japonicus TSA is also most likely contamination. Again, two Apostichopus japonicus assembled genomes based on 17 and 4 related DNA SRA experiments or 31,038 recorded proteins or 400,950 nucleotide sequences contain no reads related to putative innexin\pannexin sequences, including GHCH01013653.1. BLASTN and BLASTX search using GHCH01013653.1 sequence as a query against NCBI nr database returns crustacean sequences, such as hypothetical protein TCAL_01429 of Tigriopus californicus with E-value 7e-148 and total score 430 in BLASTX. It leads to the conclusion that the original sample was contaminated with crustacean species.

If GHCH01013653.1 is contamination then relative sequences can be present in high-throughput databases of animals with similar ecological features and no innexins in their genomes. Indeed, Megablast search using the GHCH01013653.1 as a query revealed innexin-like sequence HBWD01520106.1 in sponge TSA (Halichondria panicea, transcribed RNA sequence with E-value 7e-143 and total score 516). BLASTX of HBWD01520106.1 from Halichondria panicea TSA to NCBI proteins reveals as best hit the same crustacean innexin from Tigriopus californicus with E-value 3e-113 and total score 335. Similar sequences were also found in the brown algae Argassum vulgare (TSA: GEHA01029897) and red algae Rhodonematella subimmersa (TSA: GFTI01036537).

The sequencing data on Arbacia punctulata is small but the TSA sequence GECD01051584.1 related to putative innexin\pannexin is also probably contamination. It derives from the single Arbacia punctulata SRA experiment and is most similar to crustacean Armadillidium nasatum innexin KAB7504537.1 with E-value 1e-51 and total score 174 in BLASTX search.

Only one of 5 TSA experiments revealed sequences related to putative innexin\pannexin from Paracentrotus lividus. Paracentrotus lividus TSA sequence GIIR01058804.1 is most similar to flatworm Paragonimus heterotremus innexin KAF5405742.1 with E-value 0.0 and total score 624 in BLASTX search. The assembled Paracentrotus lividus genome based on 20 related DNA SRA experiments or 142346 recorded nucleotide sequences contain no reads related to putative innexin\pannexin sequences, including GIIR01058804.1.

Thus it appears that all 4 putative innexin\pannexin homologous proteins from 4 echinoderm species reported by Welzel and Schuster are probably cross-species contaminations. As they all derived from TSA databases, we compared the presence of putative innexin\pannexin-like sequences in echinoderm TSAs to TSA projects from other taxonomic groups. We found that as in Echinodermata, some TSA projects from taxonomic groups generally believed to have no innexins/ pannexins (Porifera, Fungi, Chlorophyta, Rhodophyta, and Phaeophyceae) contain sequences similar to innexin/pannexins. However, the number of such TSA projects does not exceed 10% of all TSA projects in examined taxonomic groups. The only exception is Porifera, where 8 out of 20 TSA projects contain innexin-like sequences (15 species of Porifera are represented and 6 species show TBLASTN hits seeded by innexin/pannexins with E-value <1e-10). Nevertheless, the fact that most Porifera TSAs still do not contain such sequences, and that innexin/pannexins are not found in Porifera annotated sequences in GenBank (501034 nucleotide and 153146 protein sequences), confirms that is still contamination. The high percentage of contaminated Porifera samples is well explained by the structure of sponges, whose bodies are the habitat of many species. On the opposite, more than 90% of TSA projects from taxons that were shown to encode innexin\pannexin proteins (Platyhelminthes, Nematoda, Annelida, Copepoda, Gastropoda, Bivalvia, Cephalopoda) contain innexin/pannexins sequences (See Fig1 and Table 1 in Supplementary materials).

**Fig. 1.**
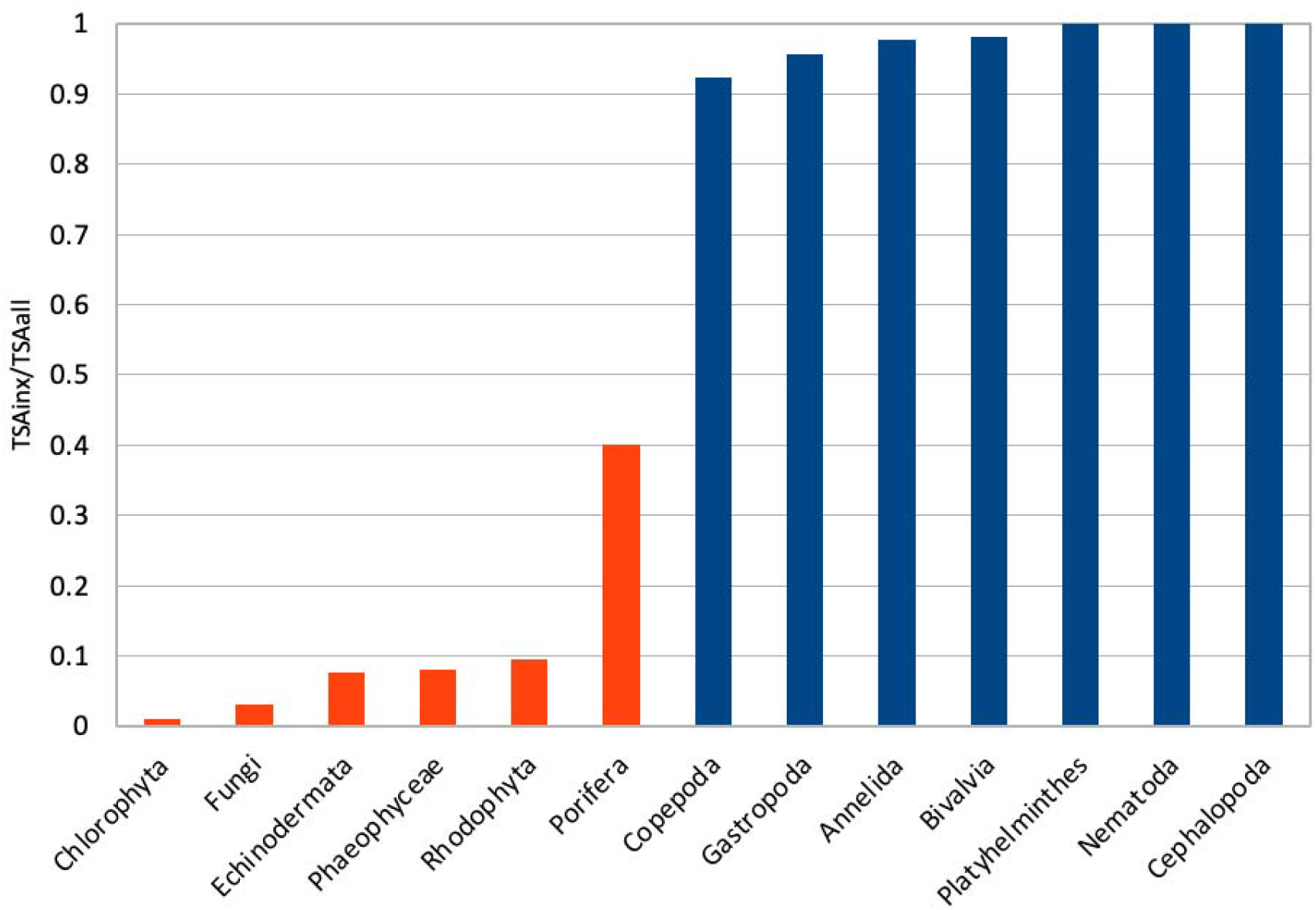
Fraction of TSAs containing innexin/pannexin -like sequences with E-value <1e-10 in different taxons. Red – taxons in which no innexins or pannexins were reported earlier. Blue – taxons with reported innexin\pannexin proteins. The taxons for comparison were selected so that the total number of available TSAs containing more than 1000 sequences was comparable to that of echinoderms (20 – 200). The taxonomic ranks are different due to the unequal representation of different groups in the databases.

Only one Echinodermata connexin-like sequence GHJZ01010392.1 was detected in the sea urchin (Mesocentrotus franciscanus) transcribed RNA sequence by Welzel and Schuster. It is most similar to chordate Geotrypetes seraphini gap junction beta-2 protein-like sequence with E-value 1e-122 and a total score of 455 in Megablast search and probably represents contamination. Connexin-like sequences were not detected in other echinoderms databases.

We therefore conclude that all connexin and innexin sequences reported by Welzel and Schuster from echinoderms are cross-species contaminations. The echinoderms most likely encode no connexin or innexin homologous sequences, which indicates that a search for a third GJ protein family is still a subject of immediate scientific interest. We also suggest that the percentage of TSAs containing the sequences of interest can be a good indicator of whether such sequences are contained in the organisms of a given taxon, or their presence is rather the result of sample contamination.

