## Supplementary materials for "Comment on “Connexins evolved after early chordates lost innexin diversity”"

**Supplementary data**

>inn sequence

MAMIATDIKGMLSVKIKPRNDTYTDQYNRIFMVKILLVTCVIMGISWFNDSVKCLVPGVNAVDGGFVSQACWIQGVYVYKELMYRSSEVGYFGIPKDMDNDGMLASGELCSTTPKFGVVNDKCKPMQKTFFLQYQWMPFLIAALSILYYLPYIGFRSANSDLISLKNTIKGGTANAEKIAKNFFDRHSNPSRNMTLRVVFNILIKVLYIVANLVAFLGLDNLLNGEFVSYGSKWVSWAKLDNAVAYDYMGKYDQPKPGNVLLPPFGYCEMYESSKDIKHSTANKHKLICELSQNILYQYSLVIVWFAIVFGIVISVIGLILLLVSYAVNMFGAKDNYGGKKIRSLTLREREYLEFIQKKNPILYREVLECLHDGNDTLPLYPQADRYNEKGAGYQLMIGSEFLDYASKLHVATYVGVEDFADKFNFLITVMVLLLCATVVTVKQYMLKPISCYIATEVGGRNLLDYVENYCWVQGTIPISYAGNVPENDEQWAKLESQKLLYYQWVPFVLGLQCILFYIPRIIWQMICYNRTGTDLQHLINLASEAAHATSDKRAETVKHLAKSLEQLLFQHREYRTGWATKVRKVCYKVCRLFYISKRLGTHLFGTYLFIKLLYLANSVGQLYLMQNFLGLNSSQYTYLGAAIATNILTGRDWQVTLVFPRVGFCVVPIRHLVGPVYATAQCALPMNMLNERIYMFLWFWVLLSAFLTAISIPMWFMRMAYEKSRTNFVRRFLRVGYDEENRYSSKDRCMVAKFTRQFLRHDGIFLLRMVAINAGELIAAEIVHKLWEIYKQKYYNRDFNVPDANEEENDSAWGGGGDSDAVRHHGPNGSAIVLMERGNALTQRPSPPSAPSHSPGLYKGKASVEDAENEEQSLRKQYLVRYRGAKADIDSPFFKLHYRTSATFCFISCLLVTANDLIGSTIDCISGTIPGNVLNTYCWIMSTFSVPSKPGGVHGEDYAYQGVEPLGDPNDRVIHAYYQWVPFVLFFQGLLFYFPHWLWKTFEDRKLDKITSGLRGRTLSLDERKDQCSILVKYVTETFHMHNFYAFKYFICDILNFINVIVQIYMINAFLGGVFMAYGSDVLYWSESASETRTDPMIDVFPRITKCNFYKYGPSGTIERHDAMCVLALNIINEKIYVFLWFWYIILAVMTSLYLLYVLAVVAVPSMRRVMVERNAKFDIKLNRDKALKQKEKMDILMRKAQMGDWFVIFLLSKNLDSILFKEFIVQLADKLKTDMPSAQQHQSTLKNGGRRQKKGQGRESSGYHHAAVVEVGAMVLQNMFSALSFLHYRKDDDFVDRLSYFYTSSFLIMMAVLVSFKQFGGRPLECWVPAQFTASWEAYTEMYCWAQNTYWVPIDQDIPVDISEREYRQISYYQWVPFFLLLQAFLYYIPCLMWRLMSDKSGIRLNDIVQMATEKENIEPDYRIRTIESLSRHIESALRYQHTATSRTQYTLHRVFKCFNMRYYESYVTGMYLATKIMYVGNILTNLVLVNKFLETDEYSIYGLGVLRDLLFGRTWIESGNFPRVTLCDFEVRVLGNNQRHSVQCVLVINIFNEKIFILIWLWFTLLFVASTLDMLYWFSISMFHRDRFRFVLRHLELTSDPDKPELFRKEKRKQVEHFLRTYLKVDGVLVLRMIALHAGVMFCTEITDALWKRYLSQHPENLIDDDSSLITFARAQSIRRKRNDSSNSDGQNHQRLGRILSHRSTGGSFRLNRTQSTNRSLNTENKTGAATGSNAAASPNTPPPVPKSACSSRFGKVKPRRTVVTINSQQNDARTSPDPIQEMSRMENVPLAVMHHLLEQSADMATALLAGEKLRELILPGSQDDKAGALAALLLQLKLELPFDRVVTIGTVLVPILLVTLVFTKNFAEEPIYCYTPHNFTRDQALYARGYCWTELRDALPGVDASLWPSLFEHKFLPYALLAFAAIMYVPALGWEFLASTRLTSELNFLLQEIDNCYHRAAEGRAPKIEKQIQSKGPGITEREKREIIENAEKEKSPEQNLFEKYLERRGRSNFLAKLYLARHVLILLLSVVPISYLCTYYATQKQNEFTCALGASPDGPVGSAGPTVRVSCKLPSVQLQRIIAGVDIVLLCFMNLIILVNLIHLFIFRKSNFIFDKLNKVGIKTRRQWRRSQFCDINILAMFCNENRDHIKSLNRLDFITNESDLMYDNVVRQLLAALAQSNHDTTPTVRDSGIQTVDPSINPAEPDGSAEPPVVKRPRKKMKWIPTSNPLPQPFKEQLAIMRVENSKTEKPKPVRRKTATDTLIAPLLDAGARAAHHYKGSGGDSGPSSAPPAASEKKHTRHFSLDVHPYILGTKKAKTEAVPPALPASRSQEGGFLSQTEECGLGLAAAPTKDAPLPEKEIPYPTEPALPGLPSGGSFHVCSPPAAPAAASLSPGSLGKADPLTILSRNATHPLLHISTLYEAREEEEGGPCAPSDMGDLLSIPPPQQILIATFEEPRTVVSTVEF

Table 1

Fraction of TSAs containing innexin/pannexin -like sequences with E-value <1e-10 in different taxons. The taxons for comparison were selected so that the total number of available TSAs containing more than 1000 sequences was comparable to that of echinoderms (20 – 200). The taxonomic ranks are different due to the unequal representation of different groups in the databases.

|  | Total number of available TSAs | TSAs containing innexin/pannexin -like sequences (E-value <1e-10) | Fraction of TSAs containing innexin/pannexin -like sequences (E-value <1e-10) |
| --- | --- | --- | --- |
| Chlorophyta | 98 | 1 | 0.01 |
| Fungi | 192 | 6 | 0.03 |
| Echinodermata | 66 | 5 | 0.08 |
| Phaeophyceae | 50 | 4 | 0.08 |
| Rhodophyta | 53 | 5 | 0.09 |
| Porifera | 20 | 8 | 0.40 |
| Copepoda | 53 | 49 | 0.92 |
| Gastropoda | 94 | 90 | 0.96 |
| Annelida | 45 | 44 | 0.98 |
| Bivalvia | 107 | 105 | 0.98 |
| Platyhelminthes | 55 | 55 | 1.00 |
| Nematoda | 89 | 89 | 1.00 |
| Cephalopoda | 33 | 33 | 1.00 |
